# Association of oropharyngeal cancer recurrence with tumor-intrinsic and immune-mediated sequelae of reduced genomic instability

**DOI:** 10.1101/2024.10.31.621311

**Authors:** Malay K. Sannigrahi, Lovely Raghav, Dominick J. Rich, Travis P. Schrank, Joseph A. Califano, John N. Lukens, Lova Sun, Iain M. Morgan, Roger B. Cohen, Alexander Lin, Xinyi Liu, Eric J. Brown, Jianxin You, Lisa Mirabello, Sambit K. Mishra, David Shimunov, Robert M Brody, Alexander T. Pearson, Phyllis A. Gimotty, Ahmed Diab, Jalal B. Jalaly, Devraj Basu

## Abstract

**Background:** Limited understanding of the biology predisposing certain human papillomavirus-related (HPV+) oropharyngeal squamous cell carcinomas (OPSCCs) to relapse impedes therapeutic personalization. We aimed to identify molecular traits that distinguish recurrence-prone tumors.

**Methods:** 50 HPV+ OPSCCs that later recurred (cases) and 50 non-recurrent controls matched for stage, therapy, and smoking history were RNA-sequenced. Groups were compared by gene set enrichment analysis, and select differences were validated by immunohistochemistry. Features discriminating groups were scored in each tumor using gene set variation analysis, and scores were evaluated for recurrence prediction ability.

**Results:** Cases downregulated pathways linked to anti-tumor immunity (FDR-adjusted p<.05) and contained fewer tumor-infiltrating lymphocytes (p<.001), including cytotoxic T-cells (p=.005). Cases also upregulated pathways related to cell division and other aspects of tumor progression. Upregulated and downregulated pathways were respectively used to define a tumor progression score (TPS) and immune suppression score (ISS) for each tumor. Correlation between TPS and ISS (r=.603, p<.001) was potentially explained by observed upregulation of DNA repair pathways in cases, which might enhance their progression directly and by limiting cytosolic DNA-induced inflammation. Accordingly, cases contained fewer double-strand breaks based on staining for phospho-RPA32 (p=.006) and γ-H2AX (p=.005) and downregulated pro-inflammatory components of the cytoplasmic DNA sensing pathway. A combined score derived from TPS and ISS optimized recurrence prediction and stratified survival in a manner generalizable to three external cohorts.

**Conclusions:** We provide novel evidence that limiting genomic instability makes tumor-intrinsic and immune-mediated contributions to HPV+ OPSCC recurrence risk, opening opportunities to detect and target this treatment-resistant biology.

## INTRODUCTION

HPV+ OPSCCs are increasing in incidence, recently surpassing cervical cancer as the most common HPV-related malignancy in the US [1, 2]. Despite relatively favorable prognosis, these cancers receive the same toxic combinations of high-dose radiation therapy with cytotoxic chemotherapy developed for more aggressive, HPV-negative head and neck cancers. In addition, a sizeable subset of patients in the US initially undergo neck dissection plus transoral robotic surgery (TORS) in an effort to avoid chemotherapy and reduce radiation doses and fields [3-5]. However, treatment of this disease is largely devoid of targeted agents and remains minimally personalized at present.

Therapeutic personalization for HPV+ OPSCCs is hindered by incomplete understanding of their distinctive biology, including the molecular underpinnings of variable therapy responses among them. The majority of patients have curative responses but can suffer severe and lasting treatment-related disabilities [6]. Averting this toxicity by de-escalating current treatments is impeded by certain tumors whose propensity to recur is not discernable from clinical features at presentation. Moreover, there is a need for agents tailored to the distinct biology of HPV+ OPSCCs, including approaches targeting vulnerabilities of the recurrence-prone subset.

The relative sensitivity of HPV+ OPSCCs to the genotoxic stress caused by radiation and cytotoxic drugs is at least partly related to the distinctive activities of HPV’s E6/E7 oncoproteins. Unrestrained G_1_-S progression caused by E6/E7 induces DNA damage by inducing high levels of replication stress [7, 8], a major source of genomic instability. Other activities of HPV oncoproteins further threaten host genome integrity by driving APOBEC3-mediated mutagenesis [9, 10] and impeding DNA repair [11-14], leaving these tumors vulnerable to excessive genomic instability produced by unresolved DNA damage.

Although genomic instability is a hallmark driver of carcinogenesis, extreme levels of it both kill tumor cells directly and enhance anti-tumor immune responses by multiple mechanisms, including activation of cytoplasmic DNA sensing pathways [15]. In many HPV+ OPSCCs, this effect may offset HPV’s multiple immune-suppressive activities [16] and contribute to their robust immune infiltrates relative to other head and neck cancers [17] and detectable activation of T-cells against viral tumor antigens [18, 19]. However, their recurrences have not proved more responsive to anti-PD-1 therapy [20], suggesting that the relapse-prone cases lack the replication stress-induced immune milieu of more typical tumors.

Prior RNA sequencing (RNAseq) studies evaluating features of HPV+ OPSCCs that recur [21-26] have been constrained by cohorts containing relatively few recurrences, limited follow-up, lack of matched controls, and/or heterogeneous therapy. In this study, we explored the biology of treatment resistance using a large patient cohort receiving TORS plus guideline-based adjuvant therapy and prolonged follow-up. To identify molecular traits with broad relevance to treatment resistance, we compared transcriptomes of tumors that recurred to those of matched controls cured by similar therapy. We describe a striking association of recurrence with tumor-intrinsic and immune-mediated effects of diminished DNA damage. These observations could guide future targeted approaches.

## METHODS

### Patients

Consecutive patients with treatment-naïve p16+ OPSCCs of tonsil and/or tongue base undergoing neck dissection plus TORS between 1/1/2007 and 12/31/2020 were identified at our institution. Charts were reviewed on 3/22/2022 (IRB #833890). Cases were defined as tumors recurring distantly and/or “in-field,” which was defined as recurrence in an “at-risk” area treated in a radiation plan. Patients with out-of-field locoregional recurrence plus distant recurrence were classified as distant recurrences for analysis. Controls without recurrence were matched as in Results, with pairing in absence of precise matches detailed in Supplementary Methods.

### DNA/RNA extraction, mRNA sequencing and processing

Formalin-fixed paraffin-embedded (FFPE) blocks were macro-dissected to maximize tumor content and sectioned into rolls for DNA/RNA extraction. DNA and RNA were isolated with the QIAamp DNA FFPE Kit (Qiagen) and Mag-Bind RNA FFPE kit (Omega Bio-tek). Sample replicates, quality control, library preparation, sequencing, alignments, and data processing are described in Supplementary Methods. Gene set enrichment analysis (GSEA) and gene set variation analysis (GSVA) are detailed in Supplementary Methods. Multivariate logistic regression was used derive coefficients provided in Supplementary Methods for weighting GSVA scores to optimize case-control segregation.

### Immunohistochemistry (IHC) analysis

Sections were IHC-stained on the Leica Bond-III^TM^ platform. Antibodies and conditions are in Supplementary Table 1. Slides were digitized using a 3DHISTITECH Panoramic Scanner. QuPath software [27] was used to define 20 tumor regions of interest per section. Cell and nuclear positivity were defined using mean intensity threshold cutoffs above background.

### Statistical methods

Characteristics of cases and controls were compared by Fisher’s exact or independent t-test. Predictive modeling combining patient characteristics with GSVA-derived scores was performed using logistic regression. Kaplan–Meier survival curves with log-rank tests were used to compare recurrence-free survival (RFS) and overall survival (OS). All tests were two-sided, with p<.05 considered significant. Multiple testing correction was performed using the Benjamini and Hochberg Procedure or Bonferroni and Sidak’s procedure. P-values for area under the ROC curve (AUC) were calculated from the z-ratio using the normal distribution. P-values in Kaplan Meier plots were calculated by log-rank test. Analyses were performed using Rv4.2 and GraphPad Prism^TM^.

## RESULTS

### Features of the case-control cohort

Case and control tumors were curated from 851 treatment-naive p16+ OPSCC patients receiving primary TORS plus neck dissection. Patient characteristics are described in Supplementary Table 2. To enrich for cases with treatment-resistant biology, we excluded tumors that recurred locoregionally outside an adjuvant radiation field, including those not receiving guideline-indicated radiation altogether. The 50 remaining cases were comprised of 36 with distant failure, 7 with locoregional failure, and 7 with both. Cases were matched 1:1 to control tumors from patients with follow-up beyond the interval to the last recurrence (53.4 months). Stage was matched using 8^th^ edition AJCC pathologic stage (T, N, and overall) rather than clinical stage because only the former stratified the total cohort for OS (p<.001) (Figure 1a). The widely cited negative prognosticator of >10 pack-years smoking and 7^th^ edition AJCC clinical ≥N2b status [6] also did not stratify OS, whereas >10 pack-years alone was significant (p=.036) (Figure 1b) and was used in matching. Lastly, receiving adjuvant radiation and systemic therapy, including the systemic agent, was used in matching. The three pathologic stage III cases were excluded due to a lack of matchable controls. Some matching imprecision was tolerated as detailed in Supplementary Methods, and one control was dual-weighted as a match for two cases for molecular comparisons due to the lack of a suitable second match, resulting in 49 unique controls. Pre-treatment tumors from metastatic lymph nodes were used preferentially for RNAseq. In four cases lacking usable nodes, primary tumor was used for both case and control. The process of curating cases and controls is depicted in Figure 1c, and Supplementary Table 3 details case-control pairs, highlighting imprecise matches. Despite imprecisions, the groups were highly similar for matched traits (Table 1), which appear representative of most HPV+ OPSCCs at presentation.

**Figure 1:**
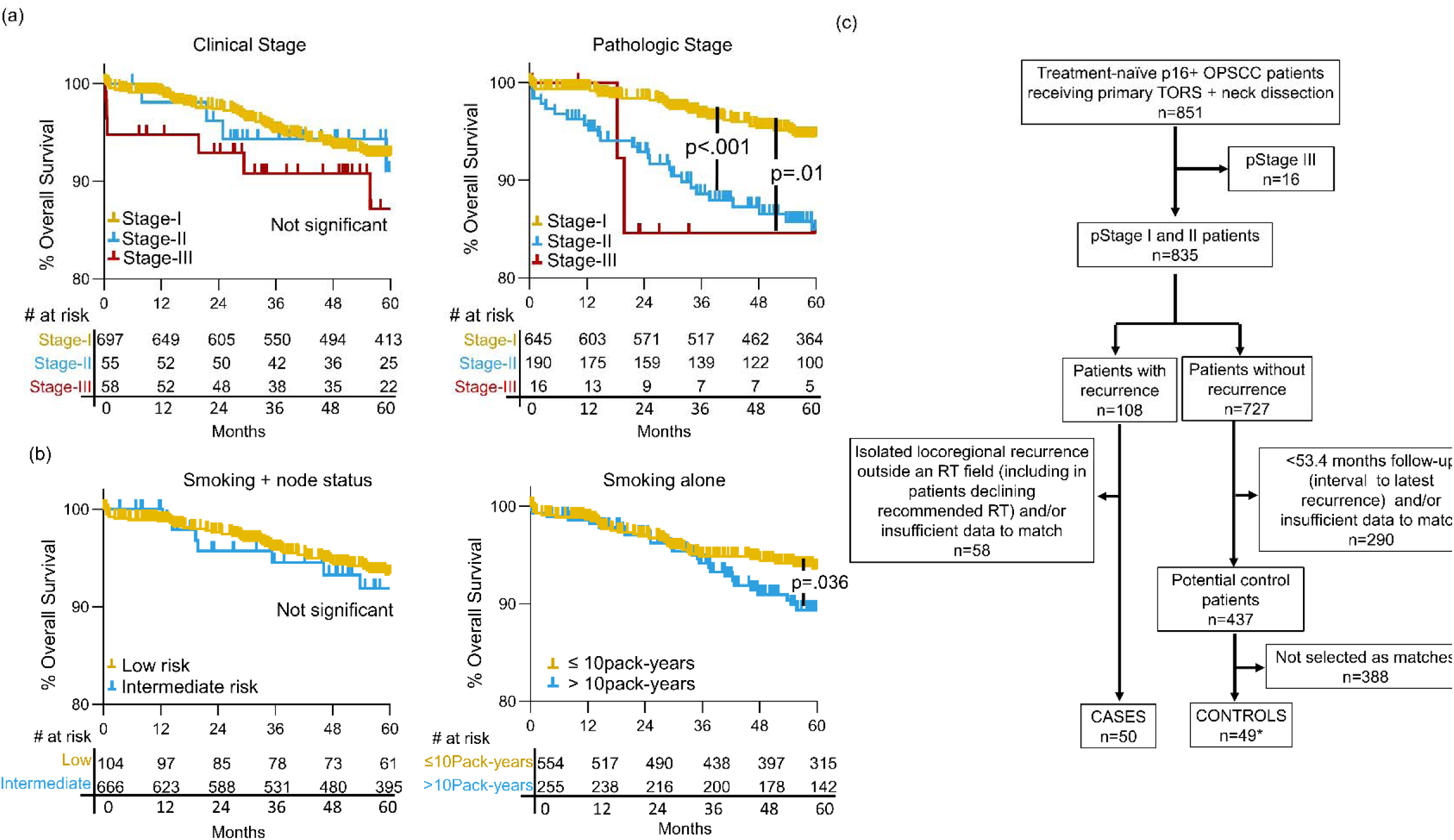
Process for selecting case and matched control tumors from the total cohort. (a) Kaplan Meier analysis of OS in total cohort by clinical (left) and pathologic (right) 8^th^ ed. AJCC stage. (b) Kaplan Meier analysis of OS in total cohort by smoking-related risk group (low risk: ≤10 pack-years and/or AJCC 7^th^ ed. <cN2b, intermediate risk: >10 pack-years and AJCC 7^th^ ed. ≥cN2b) and OS by smoking history alone (right). (c) CONSORT diagram outlining selection of cases and controls. *A single control was duplicated as a match for two cases in molecular analyses. P-values were calculated for the log-rank test.

**Table 1:** Matched features of tumors that recurred (cases) vs. non-recurrent tumors (controls)

| Variable | Categories | Case<br>(n=50) | Control<br>(n=50*) | P-value |
| --- | --- | --- | --- | --- |
| Overall pathologic stage | Stage I | 25 | 25 | 1.000 |
|  | Stage II | 25 | 25 |  |
| Pathologic T-stage | pT0 | 1 | 0 | .483 |
|  | pT1 | 11 | 15 |  |
|  | pT2 | 32 | 28 |  |
|  | pT3 | 6 | 7 |  |
| Pathologic N-stage | pN0 | 5 | 5 | .977 |
|  | pN1 | 26 | 27 |  |
|  | pN2 | 19 | 18 |  |
| Treatment | Surgery | 4 | 4 | 1.000 |
|  | Surgery + RT | 19 | 19 |  |
|  | Surgery + CRT | 27 | 27 |  |
| Systemic agent | Cisplatin | 21 | 23 | .848 |
|  | Cetuximab | 5 | 3 |  |
|  | Carboplatin/Taxol | 1 | 1 |  |
| Smoking history | ≤10 pack-years | 34 | 33 | .832 |
|  | >10 pack-years | 16 | 17 |  |

RT=radiation therapy, CRT=chemoradiation, P-values are from Fisher-exact test. Freeman-Holton extension was used for more than two categories.
\*Comparison includes duplication of one control as a match for two cases

### Cold immune microenvironments predominate in recurrence-prone tumors

Groupwise comparison of cases to controls by GSEA using the Hallmark pathways [28] (n=50) and the non-disease-specific subset of KEGG Legacy pathways [29] (n=147) identified 21 pathways downregulated in cases (FDR-adjusted p<.05). Grouping them by functional relatedness (Figure 2a) suggested suppression of innate immunity based on downregulation of Rig-I receptor signaling and interferon-α response. There were also decreases in pathways related to lymphocyte trafficking and effector function, including T-cell and B-cell receptor signaling and NK-mediated cytotoxicity. Accordingly, proinflammatory signaling pathways were downregulated, most notably interferon-γ response. Together, these results suggested attenuation of anti-tumor immunity in cases, leading us to quantify T-cells directly using IHC. Tumor-infiltrating total (CD3+) and cytotoxic (CD8+) T-cells were quantified (Figure 2b) and used to calculate Immunoscore (Figure 2c, left), a prognostic metric of anti-tumor immunity in other cancers [30]. Reductions in CD3+ (p=.036) and CD8+ (p=.005) cells in cases contributed to reduced Immunoscore (p=.006); however, Immunoscore offered limited discrimination of cases (AUC=.662, p=.005, OR=1.09, 95% CI=1.03–1.66) (Figure 2c, right). Because reduced B and NK-cell functions were also inferred from RNAseq, we quantified total tumor-infiltrating leukocytes (TILs). A marked reduction in CD45+ TIL content was observed (p<.001) (Figure 2d, left) and provided better discrimination of recurrence potential (AUC=.842, p<.001, OR=1.18, 95% CI=1.1-1.129) (Figure 2d, right). We further evaluated whether the groups are predicted to have divergent immunotherapy responses using combined positive score (CPS) [20] based on PD-L1 IHC. Cases showed lower mean CPS (p=.039) (Figure 2e, left) and more often scored <1 than ≥1 (p=.005), 1-19 (p=.032), or ≥20 (p=.013) (Figure 2e, right). These findings show that the treatment-naïve HPV+ OPSCCs predisposed to recur post-TORS-based therapy contain cold immune microenvironments predicted to be relatively refractory to immunotherapy.

**Figure 2:**
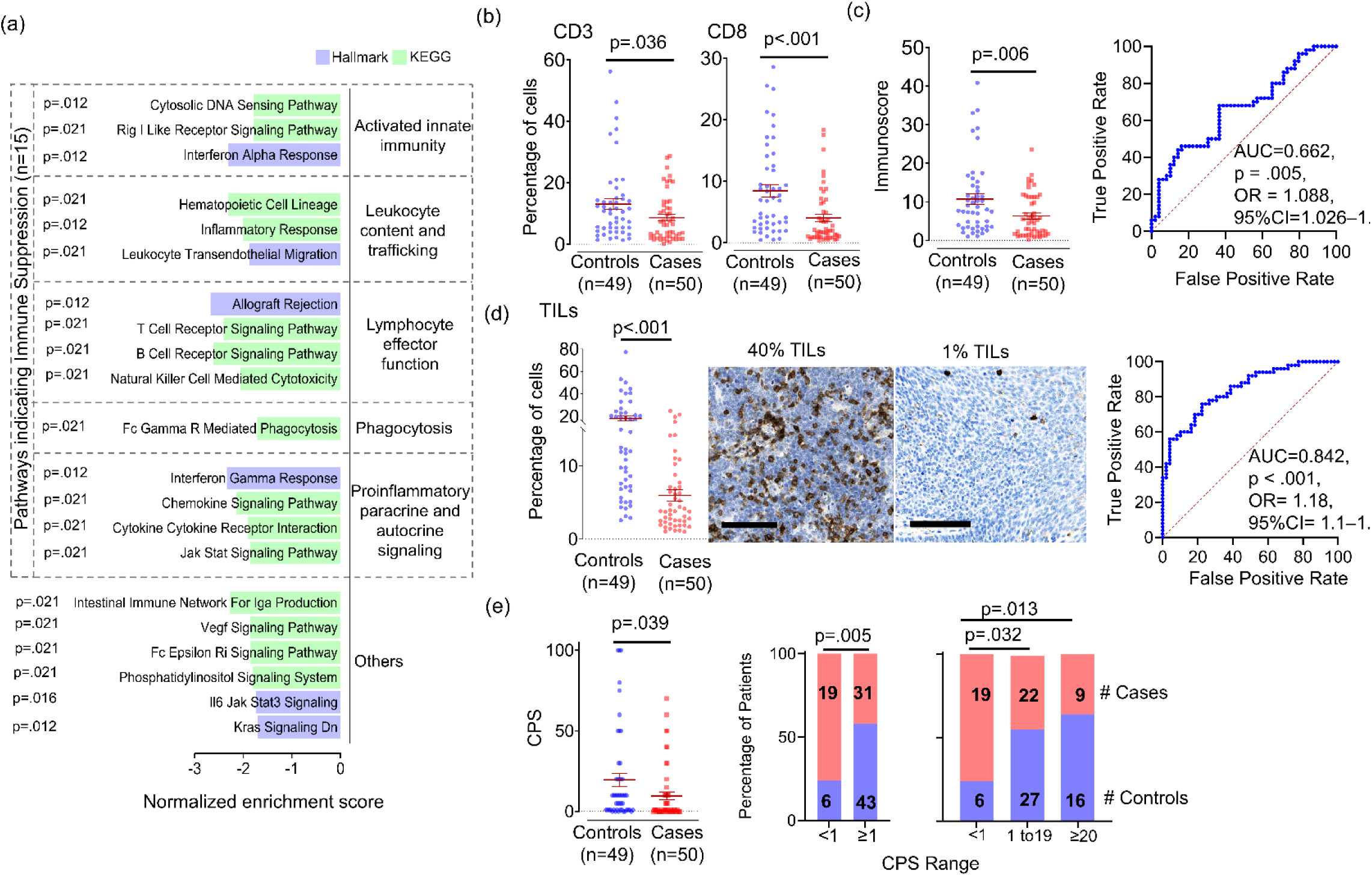
Evidence of a cold immune microenvironment in the HPV+ OPSCCs predisposed to recur. (a) Significantly downregulated pathways (FDR-adjusted p<0.05) in cases compared groupwise to controls using GSEA. Significance was determined using the Benjamini and Hochberg False Discovery method. (b) Percent CD3+ and CD8+ cells per tumor area by IHC. (c) Immunoscore derived from average of percent CD3 and CD8 cells per tumor area (left) and ROC curve for Immunoscore separating cases from controls (right). (d) Percent CD45+ cells per tumor area in cases vs. controls (left) and ROC curve for separation of cases from controls by TIL content (right). Images are 40x, bar=100µm. (e) Combined Positive Score (CPS) based on PD-L1 IHC (left) and frequency of cases and controls in CPS categories (right) with p-value calculated for two-way comparison using Fisher exact and three-way comparison using Sidak’s procedure. P-values for all other two group comparisons were calculated using unpaired t test with Welch’s correction. P-values for AUCs were calculated from the z ratio using the normal distribution. OR=odds ratio, CI= confidence interval.

### Pathways related to tumor progression and immune suppression are coordinately regulated

Additional features distinguishing the cases were evident from upregulation of 20 pathways (FDR-adjusted p<.05) in groupwise comparison to controls. Grouping by relatedness (Figure 3a) highlighted six pathways directly involved in cell division. Biosynthetic and bioenergetic demands of cell division were indirectly reflected in upregulated pathways for fatty acid synthesis, protein synthesis, oxidative metabolism, and glycolysis. These functions were consistent with the identity of the upregulated oncogenic signaling pathways (Myc targets and mTORC1 signaling), given that Myc upregulates glycolysis in cancer [31] and mTORC1 promotes translational initiation [32]. Both signaling pathways also drive biogenesis of mitochondria [33, 34], the site of oxidative metabolism and synthesis of many macromolecular precursors. Centrality of mitochondrial functions among upregulated pathways led us to quantify mitochondrial content directly using the MTCO1/β2M DNA qPCR assay [35]. The higher mitochondrial mass observed in cases (p<.001) (Figure 3b) can aid cancer progression in hostile microenvironments [36], consistent with upregulation of two invasion and metastasis-related pathways (epithelial to mesenchymal transition and ECM-receptor interaction). Together, these findings show that cases as a group upregulate transcriptional programs supporting proliferation and other aspects of tumor progression. However, it was unclear whether the tumor progression and immune suppression-related features were independent aspects of recurrence-prone biology or interrelated in individual tumors. To address this question, a tumor progression score (TPS) and immune suppression score (ISS) were developed for each tumor by GSVA of the distinct genes in the tumor progression-related and immune-suppression-related pathways, as detailed in Figure 3c. Correlation between TPS and ISS (r=.603, p<.001) (Figure 3d) indicated that transcriptional profiles directly related to tumor progression and profiles reflecting anti-tumor immune suppression are co-regulated in individual tumors.

**Figure 3:**
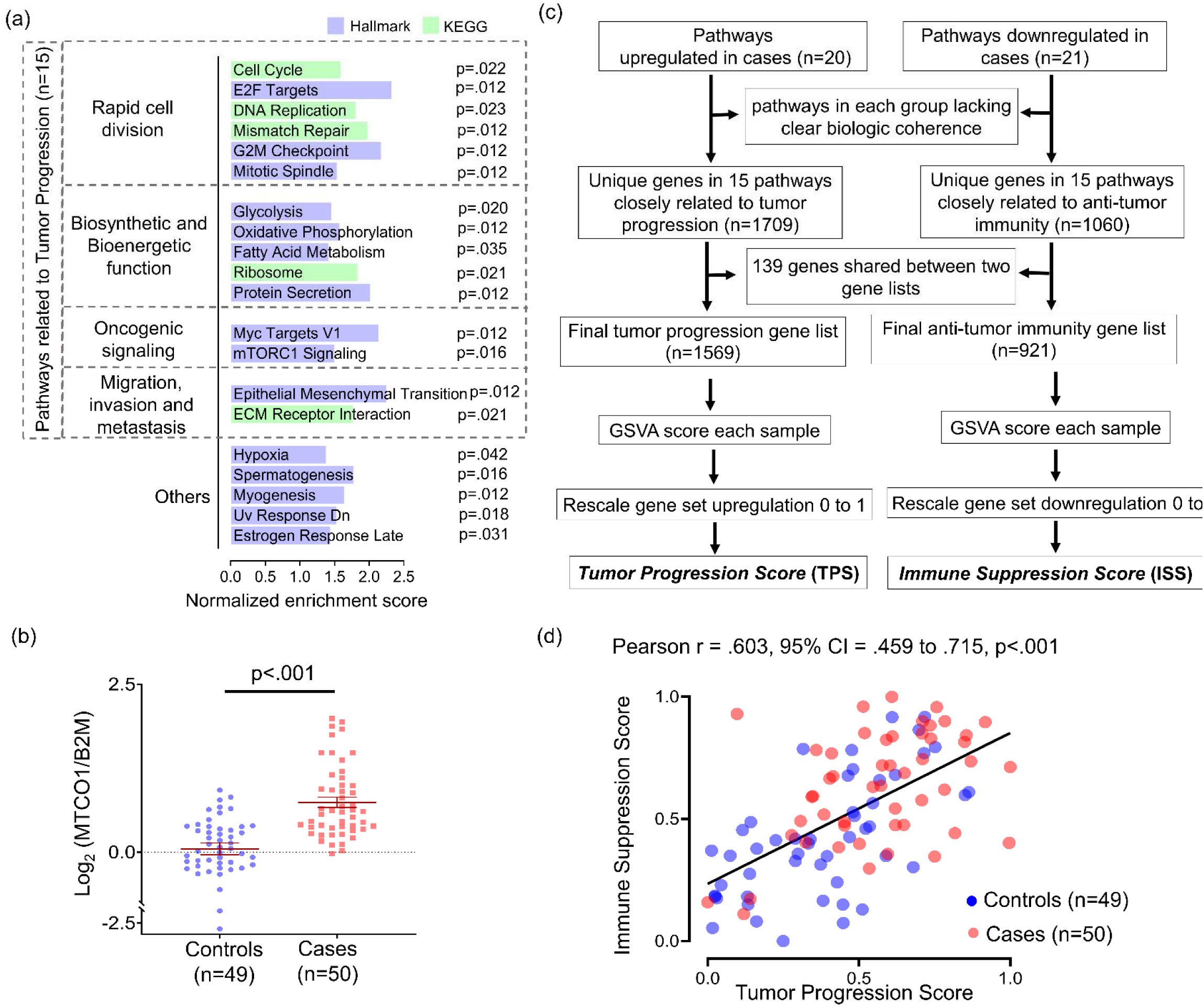
Pathways related to tumor progression and immune suppression are coordinately regulated. (a) Significantly upregulated pathways (FDR-adjusted p<0.05) in cases compared groupwise to controls using GSEA. Significance was determined using the Benjamini and Hochberg False Discovery method. (b) Comparison of mitochondrial mass expressed as MT-CO1 to β2M ratio measured using DNA qPCR. P-value was calculated using unpaired t test with Welch’s correction. (c) Flow chart illustrating process of scoring immune suppression and tumor progression. (d) Correlation plot of immune suppression score and tumor progression score across individual cases and controls. Pearson correlation coefficients were used to calculate r values, and p-values were determined by t distribution.

### Reduced DNA damage links tumor progression to immune suppression

We evaluated whether the coordinated increase in TPS and ISS in recurrence-prone tumors might arise from higher levels of HPV oncoproteins, given their multiple roles in driving de-differentiation, cell cycle progression, cell motility, and immune suppression [37]. However, neither total HPV transcripts nor alignments to individual HPV genes differed significantly between the 46 (92%) cases and 44 (90%) controls containing HPV16 (Figure 4a). We also considered whether recurrence-prone cases more effectively limit the genomic instability that HPV oncoproteins induce via unchecked G1-S progression [7] and DDR inhibition [11-14], as relatively intact DNA repair would jointly facilitate efficient cell division and prevent the micronuclei formation that triggers inflammation via cytoplasmic DNA sensing [38]. To test this hypothesis, we examined the four DDR signaling pathways using their annotations in the Protein Interaction Database [39]: ATR signaling, ATM signaling, DNA-PK signaling, and the Fanconi Anemia (FA) pathway. GSEA showed significant ATM and FA pathway enrichment in cases (FDR-adjusted p=.002) (Figure 4b), suggesting that relatively intact double-strand break (DSB) repair in S-phase limits genomic instability. Direct evidence was provided by IHC for the canonical DSB markers S139-phosphorylated H2AX (γ-H2AX) and S4/S8-phosphorylated RPA32 (pRPA32) [40, 41]. A decrease in pan-nuclear γ-H2AX (p=.006) and pRPA32 (p=.005) (Figure 4c) in tumor cells of cases was consistent with their downregulation of the downstream KEGG Cytosolic DNA Sensing Pathway (Figure 2a). Scoring each tumor by GSVA for all 34 detected transcripts in this pathway further highlighted differences between cases and controls (p<.001) (Figure 4d). In groupwise comparison, cases broadly downregulated factors with pro-inflammatory function in this pathway, including the cytosolic DNA sensing master regulator STING1 (p=0.035), its downstream effectors IRF3 (p=.007), IRF7 (p=.011), and NFKB1(p=.003), as well as interferon-stimulated genes ZBP1 (p=.001) and CCL5 (p=.003) (Figure 4e). Together, these data indicate that both tumor-intrinsic and immune-mediated effects of reduced DNA damage are associated with recurrence.

**Figure 4:**
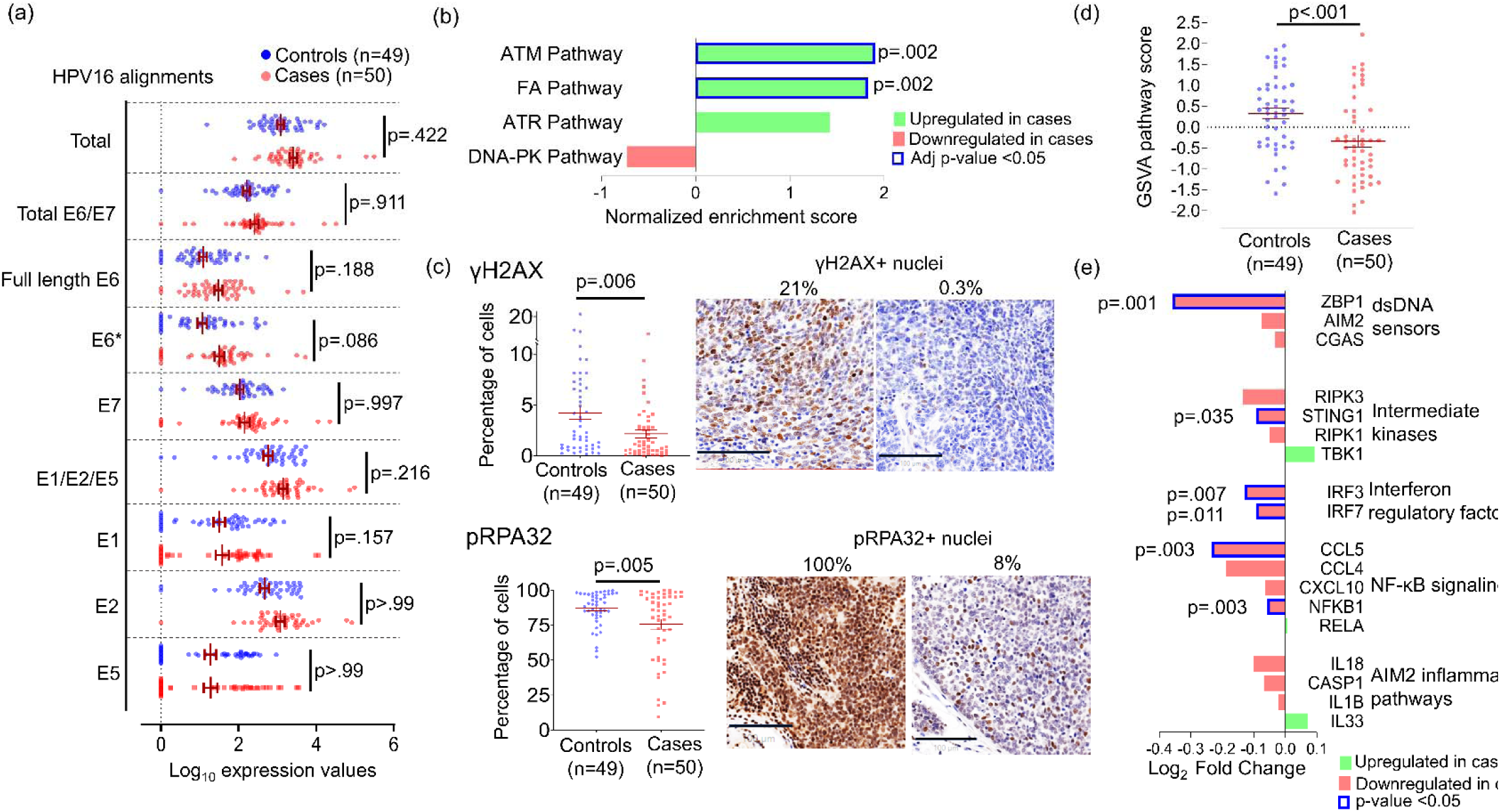
Intact DNA repair and reduced double strand breaks in HPV+ OPSCCs predisposed to ecur. (a) Normalized HPV16 transcript alignments in cases and controls. E6* indicates all spliced E6 forms. Adjusted p-values were calculated using Bonferroni and Sidak’s procedure. (b) DNA repair signaling pathways in the Protein Interaction Database compared between cases and controls using GSEA. P-values were calculated using the Benjamini and Hochberg False Discovery method. (c) Percent γH2AX-positive nuclei (top) and percent pRPA32-positive nuclei (bottom) per tumor area by IHC. Images are 40x with bar=100µm. P-va ues were calculated using unpaired t test with Welch’s correction. (d) Comparison of GSVA scores for the KEGG cytosolic DNA pathway transcripts between cases and controls. P-value was calculated using unpaired t test with Welch’s correction. (e) Expression of each gene in the KEGG cytosolic DNA Sensing Pathway with pro-inflammatory functions compared between case and control groups. P-value for each comparison was calculated using unpaired t test.

### Combining TPS with ISS optimizes recurrence prediction and stratifies survival

We evaluated whether jointly considering the features underlying TPS and ISS along with other clinical traits optimizes recurrence prediction and stratification of survival outcomes. The AUCs segregating cases from controls were 0.713 using TPS and 0.735 using ISS. Adding TPS and ISS after weighting each by coefficients derived from multivariate logistic regression yielded an improved combined score with AUC of 0.757 and a favorable odds ratio (p<0.001, OR=4.48, 95% CI=2.23-9.85) (Figure 5a). We then tested whether clinical characteristics not matched upfront enhanced recurrence prediction when considered together with combined score (Table 2). Comparing unmatched traits between cases and controls identified only lymphovascular space invasion (LVSI) and positive node number as significant. Evaluating these two features with combined score in a multivariate model showed that only LVSI maintained significance, albeit marginally (p=.046), supporting the score’s utility over clinical and pathologic information in defining recurrence-prone biology. The combined score’s ability to stratify outcomes was retained in the post-recurrence context, where a Youden index cut-point divided the 50 cases into equal-sized groups with divergent OS from time of recurrence (p=.007) (Figure 5b).

**Figure 5:**
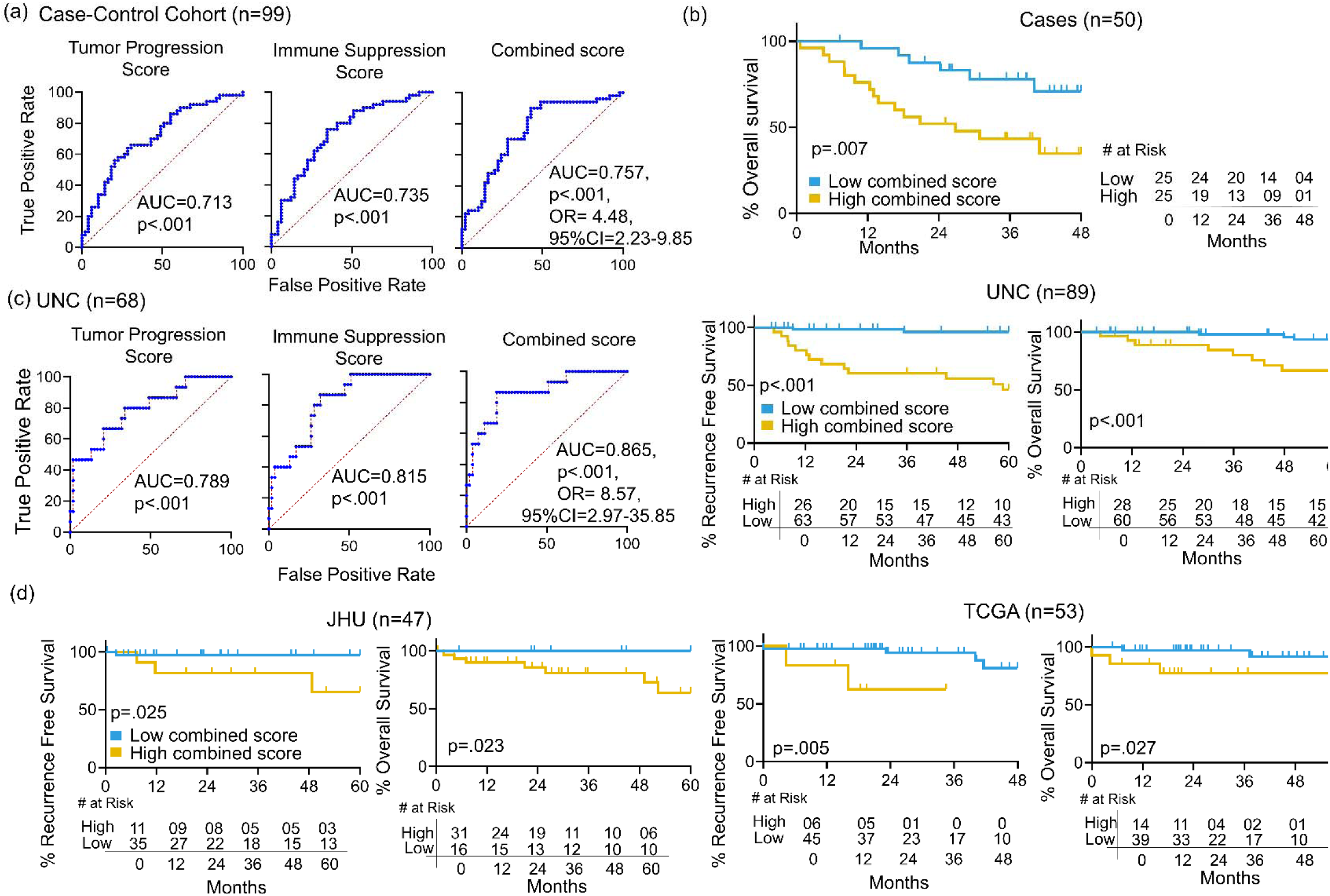
Prediction of recurrence and survival stratification using combined score. (a) ROC curves for predicting recurrence (cases vs. controls) using TPS, ISS, and Combined Score. (b) Kaplan Meier analysis using a Youden index cut-point for combined score to stratify OS of the 50 cases from time of recurrence. (c) ROC curves for TPS, ISS and Combined Score in the UNC cohort (left). Kaplan Meier survival curves for RFS and OS stratified using a Youden index cut-point for combined score in the UNC cohort (right). (d) Kaplan Meier survival curves for RFS and OS stratified using a Youden index cut-point for combined score in the JHU cohort (left) and the TCGA cohort (right). OR=odds ratio, CI= confidence interval.

**Table 2:** Contribution of unmatched features to recurrence prediction using logistic regression

| Features | Univariate model |  |  | Multivariate Model |  |  |
| --- | --- | --- | --- | --- | --- | --- |
|  | OR | 95% CI | P-value | OR | 95% CI | P-value |
| Combined Score | 4.37 | 2.185-9.606 | <.001 | 3.91 | 1.921-8.713 | <.001 |
| LVSI | 3.53 | 1.483-8.931 | .005 | 2.75 | 1.028-7.713 | .047 |
| Positive Nodes | 1.09 | 1.011-1.211 | .041 | 1.04 | 0.952-1.164 | .373 |
| Age | 1.04 | 1.001-1.105 | .051 |  |  |  |
| ENE | 1.32 | 0.599-2.963 | .487 |  |  |  |
| Sex | 1.50 | 0.444-5.412 | .516 |  |  |  |
| Level 4/5 nodes* | 1.89 | 0.717-5.276 | .204 |  |  |  |
| Race | 0.35 | 0.023-1.515 | .261 |  |  |  |
| Body mass index | 0.98 | 0.906-1.068 | .728 |  |  |  |
| Margin | 0.44 | 0.112-1.525 | .213 |  |  |  |

OR=odds ratio, CI= confidence interval, LVSI= lymphovascular space invasion, ENE= extranodal extension
\*indicates pathologically positive lymph nodes identified in neck levels 4 and/or 5

Generalizability was further tested in three independent HPV+ OPSCC cohorts, UNC [22], JHU [42], and TCGA [43], which were the only datasets identified with raw RNAseq data accessible for generating GSVA-based scores by methods identical to those used for our cohort. Relative to our case-control cohort, each validation cohort contained fewer tumors, far fewer recurrences, and a mix of patients with surgical vs. nonsurgical therapy (Supplementary Table 4). Only UNC contained sufficient cases (n=18) and controls with adequate follow-up (n=53) for testing recurrence classification by TPS and ISS, which yielded AUCs of 0.789 and 0.815 respectively. A combined score generated by the same formula used for our cohort improved AUC to 0.865 (p<.001, OR=8.57, 95% CI=2.97-35.85), (Figure 5c, left). Accordingly, Youden index-based cut-points for combined score in the full UNC cohort stratified RFS (p<.001) and OS (p<.001) (Figure 5c, right). Despite few recurrence events in JHU (n=4) and TCGA (n=6), identically generated combined scores and cut-points stratified RFS and OS in both (Figure 5d). These findings demonstrate that the predictive molecular traits jointly captured by TPS and ISS in our cohort are generalizable to more heterogeneous populations and also stratify survival post-recurrence.

## DISCUSSION

This study uncovers a network of molecular traits distinguishing recurrence-prone HPV+ OPSCCs that have clinical and pathologic features typical of most HPV+ OPSCCs at presentation. Despite analyzing tumors receiving TORS-based therapy, we detected transcriptional underpinnings of therapy resistance that also proved relevant to non-surgical treatment. Our study design partly overcomes limitations of prior RNAseq studies that contained fewer recurrences [21-26] and provided narrower insights into recurrence-prone biology. Markedly lower immune-related gene expression and TIL content in tumors that recurred adds to growing evidence for the association of HPV+ OPSCC recurrence risk with cold immune microenvironments [17, 24, 44-46]. We provide novel evidence that this unfavorable immune milieu is linked to reduced DNA damage, which also supports cancer progression via tumor cell-intrinsic effects. Our findings open opportunities to develop molecular biomarkers incorporating these multiple interrelated dimensions of high-risk biology and to target this network of effects using inhibitors of DNA repair that are emerging as anti-cancer agents [47].

Our results highlight clinical relevance for the DNA damage potently induced by HPV oncoproteins. Unchecked S-phase entry driven by E7 stalls replication forks and increases replication origin firing, eventually leading to DSBs [48]. These effects may be exacerbated by attenuation of DDR pathways via multiple E6/E7-induced mechanisms [49-51]. However, HPV’s effects on DNA repair seem paradoxical since rapid S-phase entry also induces expression of components of the ATM, ATR, and FA signaling pathways [52], which orchestrate faithful DNA repair to maintain genomic integrity. This effect during HPV’s normal life cycle may facilitate viral episome replication to high copy number [53, 54] while diverting these resources from preserving host genome integrity. The diverse effects of HPV on host DNA repair make it challenging to speculate on factors limiting DNA damage in recurrence-prone tumors, particularly given lack of clear differences in HPV oncogene expression in this study.

Adverse molecular traits in our analysis that confirm some prior observations may be of particular interest for biomarker development. Low NF-κB-mediated pro-inflammatory signaling has been linked to worse outcomes through downregulation as part of a co-expressed set of 203 genes [22]. Although this expression module was not differentially expressed in our tumors (not shown), the cases had reduced expression of pro-inflammatory NF-κB pathway components, and downregulated cytoplasmic DNA sensing provides a reasonable explanation for this effect. High mitochondrial mass is another adverse feature [26, 55] that was validated here.

Mitochondria provide critical precursors for DNA nucleotide synthesis [56] and thus help prevent replication stress-induced deoxynucleotide insufficiency in S phase [57]. However they also support antioxidant functions [58] that promote radioresistance, as demonstrated by us in HPV+ OPSCCs [26]. Thus, high mitochondrial mass offers a potential unifying basis for many transcriptional signals observed here. Rank-based quantification of these numerous signals with GSVA optimized prediction in this study; however, GSVA is ill-suited to clinical application, where optimal combinations of discrete markers remain to be defined.

Our analysis also suggests new therapeutic options for tumors with innate therapy resistance. Despite high overall TIL content in HPV+ OPSCCs, an immune-cold microenvironment in the tumors predestined to relapse likely contributes to their modest anti-PD-1 antibody responses after recurrence [20, 59]. There is ongoing interest in reversing anti-tumor immune suppression using neoadjuvant therapy based on evidence that the standard definitive treatments for head and neck cancers ablate the tumor antigens, T-cells, and lymphatic networks that mediate immunotherapy responses [60, 61]. Pharmacologically targeting DDR signaling is an appealing strategy for tumors where reduced DNA damage contributes both immune-mediated and tumor cell-intrinsic dimensions to therapy resistance. Emerging inhibitors targeting DDR signaling through ATR [62], its downstream effector CHEK1 [63], and WEE1 [64, 65] in combination with immunotherapy offer appealing strategies to prevent relapse of tumors with treatment-refractory features.

## Data availability

(repository information pending)

## Author contributions

Malay K. Sannigrahi, PhD (Conceptualization, Data curation, Formal analysis, Investigation, Methodology, Validation, Visualization, Writing—original draft, Writing—review & editing), Lovely Raghav, PhD (Conceptualization; Data curation; Formal analysis; Project administration; Software; Supervision; Validation; Visualization; Writing—original draft), Dominick J. Rich, BSc (Conceptualization; Data curation; Formal analysis; Investigation; Methodology; Visualization), Travis P. Schrank, MD, PhD (Data curation; Investigation; Methodology; Resources; Software; Writing—review & editing), Joseph A. Califano, MD (Data curation; Resources), John N. Lukens, MD (Conceptualization; Writing—review & editing), Lova Sun, MD (Conceptualization; Writing—review & editing), Iain M. Morgan, PhD (Writing—review & editing), Roger B. Cohen, MD (Conceptualization; Writing—review & editing), Alexander Lin, MD (Data curation; Formal analysis), Xinyi Liu, PhD (Data curation; Formal analysis), Eric J. Brown, PhD (Conceptualization; Writing—review & editing), Jianxin You, PhD (Funding acquisition, Writing—review & editing) Lisa Mirabello, PhD (Data curation; Formal analysis), Sambit K. Mishra, PhD (Data curation; Formal analysis), David Shimunov, MD (Data curation; Formal analysis), Robert M Brody, MD (Data curation; Formal analysis), Alexander T Pearson, MD, PhD (Conceptualization; Formal analysis; Investigation; Methodology), Phyllis A. Gimotty, PhD (Conceptualization; Formal analysis; Funding acquisition; Methodology; Project administration; Resources; Software; Supervision; Validation; Writing—review & editing), Ahmed Diab, PhD (Conceptualization; Investigation; Methodology; Supervision; Writing—review & editing), Jalal B. Jalaly, MBBS (Data curation; Formal analysis; Funding acquisition; Investigation; Methodology; Project administration; Resources; Supervision), Devraj Basu, MD, PhD (Conceptualization; Data curation; Formal analysis; Funding acquisition; Investigation; Methodology; Project administration; Resources; Supervision; Validation; Visualization; Writing—original draft; Writing— review & editing)

## Acknowledgments

We wish to thank the numerous patients with head and neck cancer whose surgical tumor specimens contributed to the insights in this study. A part of this study was presented in 2024 American Society of Clinical Oncology (ASCO) Annual Meeting May 31, 2024 - Jun 04, 2024, and published as conference abstract (DOI: 10.1200/JCO.2024.42.16_suppl.6055).

