## Supplementary Files for "Association of oropharyngeal cancer recurrence with tumor-intrinsic and immune-mediated sequelae of reduced genomic instability"

**Supplementary Table 1: Antibodies and staining conditions**

| Protein | Clone | Supplier | Catalog# | Species | Dilution | HIER* | Protocol ID† |
|---|---|---|---|---|---|---|---|
| CD3 | LN10 | Leica | PA0553 | Mouse | pre-diluted | ER2/20 min | HUP Refine |
| CD8 | c8/144b | Dako | M7103 | Mouse | 1:40 | ER1/20 min | HUP Refine |
| CD45 (LCA) | 2B11 + PD7/26 | Dako | M0701 | Mouse | 1:200 | ER1/10 min | HUP Refine |
| gH2AX | 20E3 | Cell Signaling | 9718S | Rabbit | 1:50 | ER2/20 min | HUP 60/20 |
| PDL1 | E1J2J | Cell Signaling | 15165BF | Rabbit | 1:500 | ER2/20 min | HUP 60/20 |
| S4/8 pRPA32 | (polyclonal) | Abcam | ab87277 | Rabbit | 1:100 | ER2/20 min | HUP Refine |

***HIER**: Heat-induced epitope retrieval method (detailed in Supplementary Methods)
**†Protocol ID:** The two different IHC staining protocols are detailed in Supplementary Methods.

**Supplementary Table 2: Characteristics of total cohort (n=851)**

| Variable | Categories | Number | Percent |
|---|---|---|---|
| Overall pathologic stage (8th edition AJCC) | Stage I | 645 | 75.8% |
| | Stage II | 190 | 22.3% |
| | Stage III | 16 | 1.9% |
| Pathologic T-stage | pT0 | 3 | 0.4% |
| | pT1 | 350 | 41.1% |
| | pT2 | 412 | 48.4% |
| | pT3 | 69 | 8.1% |
| | pT4 | 17 | 2.0% |
| Pathologic N-stage | pN0 | 125 | 14.7% |
| | pN1 | 590 | 69.3% |
| | pN2 | 133 | 15.6% |
| | pNx | 3 | 0.4% |
| Treatment | Surgery only | 187 | 22.0% |
| | Surgery + radiotherapy | 311 | 36.6% |
| | Surgery + chemoradiotherapy | 250 | 29.4% |
| | Unknown | 103 | 12.1% |
| Smoking history | ≤10 pack-years | 554 | 65.1% |
| | >10 pack-years | 255 | 30.0% |
| | Unknown | 42 | 4.9% |
| Overall clinical stage (8th edition AJCC) | Stage I | 697 | 81.9% |
| | Stage II | 55 | 6.5% |
| | Stage III | 58 | 6.8% |
| | Unknown | 41 | 4.8% |
| Clinical T-stage | cT1 | 213 | 25.0% |
| | cT2 | 382 | 44.9% |
| | cT3 | 32 | 3.8% |
| | cT4 | 47 | 5.5% |
| | cTx | 109 | 12.8% |
| | cT0 | 9 | 1.1% |
| | Unknown | 59 | 6.9% |
| Clinical N-stage | cN0 | 121 | 14.2% |
| | cN1 | 634 | 74.5% |
| | cN2 | 30 | 3.5% |
| | cN3 | 5 | 0.6% |
| | Unknown | 61 | 7.2% |
| Primary tumor site | Tonsil | 459 | 53.9% |
| | Tongue base | 317 | 37.3% |
| | Overlap | 62 | 7.3% |
| | Synchronous | 5 | 0.6% |

**(continued)**

| | | | |
|---|---|---|---|
| | Unknown primary | 8 | 0.9% |
| Pathologic level IV/V nodes | No | 744 | 87.4% |
| | Yes | 77 | 9.1% |
| | Unknown | 30 | 3.5% |
| Margin status | Negative | 610 | 71.7% |
| | In-situ or Close (<2mm) | 160 | 18.8% |
| | Positive | 57 | 6.7% |
| | Unknown | 24 | 2.8% |
| Lymphovascular space Invasion | No | 559 | 65.7% |
| | Yes | 256 | 30.1% |
| | Unknown | 36 | 4.2% |
| Perineural invasion | No | 689 | 81.0% |
| | Yes | 125 | 14.7% |
| | Unknown | 37 | 4.4% |
| Pathologic extranodal extension | No | 599 | 70.4% |
| | Yes | 228 | 26.8% |
| | Unknown | 24 | 2.8% |
| # positive nodes | Mean | 2.62 | |
| | Range | 0-48 | |
| Age | Median | 60 | |
| | Range | 32-89 | |
| Sex | Male | 741 | 87.1% |
| | Female | 110 | 12.9% |
| Race | White | 785 | 92.2% |
| | Non-white | 62 | 7.3% |
| | Unknown | 4 | 0.5% |
| Charlson comorbidity index | 0 | 562 | 66.0% |
| | ≥1 | 149 | 17.5% |
| | Unknown | 140 | 16.5% |

**Supplementary Table 3: Characteristics of matched cases and controls highlighting imperfectly matched traits (red text)**

| Pair | Recurrence | | | | | | Control | | | | | |
|---|---|---|---|---|---|---|---|---|---|---|---|---|
| number | Smoking | pT | pN | pOverall | Treatment | Systemic agent | Smoking | pT | pN | pOverall | Treatment | Systemic agent |
| 1 | ≤10pk-yr | pT1 | pN2 | 2 | Surgery + CRT | Cisplatin | ≤10pk-yr | pT1 | pN2 | 2 | Surgery + CRT | Cisplatin |
| 2 | ≤10pk-yr | pT3 | pN1 | 2 | Surgery + CRT | Cisplatin | ≤10pk-yr | pT3 | pN1 | 2 | Surgery + CRT | Cisplatin |
| 3 | >10pk-yr | pT2 | pN2 | 2 | Surgery + CRT | Cisplatin | >10pk-yr | pT2 | pN2 | 2 | Surgery + CRT | Cisplatin |
| 4 | ≤10pk-yr | pT2 | pN1 | 1 | Surgery + CRT | Cisplatin | ≤10pk-yr | pT2 | pN1 | 1 | Surgery + CRT | Cisplatin |
| 5 | ≤10pk-yr | pT2 | pN1 | 1 | Surgery + RT | | ≤10pk-yr | pT2 | pN1 | 1 | Surgery + RT | |
| 6 | ≤10pk-yr | pT1 | pN1 | 1 | Surgery + RT | | ≤10pk-yr | pT1 | pN1 | 1 | Surgery + RT | |
| 7 | ≤10pk-yr | pT1 | pN1 | 1 | Surgery + RT | | ≤10pk-yr | pT1 | pN1 | 1 | Surgery + RT | |
| 8 | ≤10pk-yr | pT2 | pN1 | 1 | Surgery + CRT | Cisplatin | ≤10pk-yr | pT2 | pN1 | 1 | Surgery + CRT | Cisplatin |
| 9 | ≤10pk-yr | pT2 | pN2 | 2 | Surgery + CRT | Carboplatin/paclitaxel | ≤10pk-yr | pT2 | pN2 | 2 | Surgery + CRT | Carboplatin/paclitaxel |
| 10 | ≤10pk-yr | pT2 | pN2 | 2 | Surgery + CRT | Cisplatin | ≤10pk-yr | pT2 | pN2 | 2 | Surgery + CRT | Cisplatin |
| 11 | >10pk-yr | pT2 | pN2 | 2 | Surgery + CRT | Cetuximab | >10pk-yr | pT2 | pN2 | 2 | Surgery + CRT | Cetuximab |
| 12* | ≤10pk-yr | pT3 | pN0 | 2 | Surgery + RT | | ≤10pk-yr | pT3 | pN0 | 2 | Surgery + RT | |
| 14 | >10pk-yr | pT2 | pN1 | 1 | Surgery Only | | >10pk-yr | pT2 | pN1 | 1 | Surgery Only | |
| 15 | >10pk-yr | pT3 | pN1 | 2 | Surgery + CRT | Cisplatin | >10pk-yr | pT3 | pN1 | 2 | Surgery + CRT | Cisplatin |
| 16 | >10pk-yr | pT0 | pN2 | 2 | Surgery + CRT | Cetuximab | >10pk-yr | pT2 | pN2 | 2 | Surgery + CRT | Cisplatin |
| 18 | ≤10pk-yr | pT1 | pN2 | 2 | Surgery + CRT | Cetuximab | ≤10pk-yr | pT1 | pN2 | 2 | Surgery + CRT | Cisplatin |
| 19 | ≤10pk-yr | pT2 | pN2 | 2 | Surgery + CRT | Cisplatin | ≤10pk-yr | pT2 | pN2 | 2 | Surgery + CRT | Cisplatin |
| 20 | ≤10pk-yr | pT2 | pN2 | 2 | Surgery + CRT | Cisplatin | ≤10pk-yr | pT2 | pN2 | 2 | Surgery + CRT | Cisplatin |
| 21 | ≤10pk-yr | pT2 | pN2 | 2 | Surgery + CRT | Cisplatin | ≤10pk-yr | pT2 | pN2 | 2 | Surgery + CRT | Cisplatin |
| 22 | ≤10pk-yr | pT3 | pN1 | 2 | Surgery + CRT | Cisplatin | ≤10pk-yr | pT3 | pN1 | 2 | Surgery + CRT | Cisplatin |
| 23 | ≤10pk-yr | pT1 | pN2 | 2 | Surgery + CRT | Cisplatin | ≤10pk-yr | pT1 | pN2 | 2 | Surgery + CRT | Cisplatin |
| 24 | >10pk-yr | pT2 | pN2 | 2 | Surgery + CRT | Cisplatin | >10pk-yr | pT1 | pN2 | 2 | Surgery + CRT | Cisplatin |
| 25 | >10pk-yr | pT1 | pN2 | 2 | Surgery + CRT | Cetuximab | >10pk-yr | pT1 | pN2 | 2 | Surgery + CRT | Cetuximab |
| 26* | ≤10pk-yr | pT3 | pN0 | 2 | Surgery + RT | | ≤10pk-yr | pT3 | pN0 | 2 | Surgery + RT | |
| 27 | ≤10pk-yr | pT1 | pN2 | 2 | Surgery + RT | | ≤10pk-yr | pT1 | pN2 | 2 | Surgery + RT | |
| 28 | ≤10pk-yr | pT2 | pN2 | 2 | Surgery Only | | ≤10pk-yr | pT3 | pN1 | 2 | Surgery Only | |
| 29† | >10pk-yr | pT2 | pN2 | 2 | Surgery + RT | | >10pk-yr | pT1 | pN2 | 2 | Surgery + RT | |
| 30 | ≤10pk-yr | pT2 | pN1 | 1 | Surgery + CRT | Cisplatin | ≤10pk-yr | pT2 | pN1 | 1 | Surgery + CRT | Cisplatin |
| 31 | ≤10pk-yr | pT1 | pN1 | 1 | Surgery + CRT | Cisplatin | ≤10pk-yr | pT1 | pN1 | 1 | Surgery + CRT | Cisplatin |
| 32 | ≤10pk-yr | pT2 | pN1 | 1 | Surgery + CRT | Cetuximab | ≤10pk-yr | pT2 | pN1 | 1 | Surgery + CRT | Cetuximab |
| 33 | ≤10pk-yr | pT2 | pN1 | 1 | Surgery + CRT | Cisplatin | ≤10pk-yr | pT2 | pN1 | 1 | Surgery + CRT | Cisplatin |
| 34 | >10pk-yr | pT2 | pN1 | 1 | Surgery + CRT | Cisplatin | >10pk-yr | pT2 | pN1 | 1 | Surgery + CRT | Cisplatin |

**(continued)**

| | | | | | | | | | | | | |
|---|---|---|---|---|---|---|---|---|---|---|---|---|
| 35 | >10pk-yr | pT2 | pN1 | 1 | Surgery + CRT | Cisplatin | >10pk-yr | pT2 | pN1 | 1 | Surgery + CRT | Cisplatin |
| 36 | ≤10pk-yr | pT2 | pN1 | 1 | Surgery + RT | | ≤10pk-yr | pT2 | pN1 | 1 | Surgery + RT | |
| 37 | ≤10pk-yr | pT2 | pN1 | 1 | Surgery + RT | | ≤10pk-yr | pT1 | pN1 | 1 | Surgery + RT | |
| 38 | ≤10pk-yr | pT2 | pN1 | 1 | Surgery + RT | | ≤10pk-yr | pT2 | pN1 | 1 | Surgery + RT | |
| 39 | ≤10pk-yr | pT2 | pN1 | 1 | Surgery + RT | | ≤10pk-yr | pT2 | pN1 | 1 | Surgery + RT | |
| 40 | ≤10pk-yr | pT2 | pN1 | 1 | Surgery + RT | | ≤10pk-yr | pT1 | pN1 | 1 | Surgery + RT | |
| 41* | ≤10pk-yr | pT2 | pN0 | 1 | Surgery + RT | | ≤10pk-yr | pT2 | pN0 | 1 | Surgery + RT | |
| 42 | ≤10pk-yr | pT1 | pN1 | 1 | Surgery + RT | | ≤10pk-yr | pT1 | pN1 | 1 | Surgery + RT | |
| 43 | ≤10pk-yr | pT2 | pN1 | 1 | Surgery + RT | | ≤10pk-yr | pT2 | pN1 | 1 | Surgery + RT | |
| 44 | ≤10pk-yr | pT2 | pN1 | 1 | Surgery + RT | | ≤10pk-yr | pT2 | pN1 | 1 | Surgery + RT | |
| 45 | >10pk-yr | pT2 | pN1 | 1 | Surgery + RT | | >10pk-yr | pT2 | pN1 | 1 | Surgery + RT | |
| 46 | >10pk-yr | pT2 | pN0 | 1 | Surgery Only | | >10pk-yr | pT2 | pN0 | 1 | Surgery Only | |
| 48 | >10pk-yr | pT1 | pN1 | 1 | Surgery Only | | >10pk-yr | pT1 | pN1 | 1 | Surgery Only | |
| 49† | >10pk-yr | pT1 | pN2 | 2 | Surgery + RT | | >10pk-yr | pT1 | pN2 | 2 | Surgery + RT | |
| 50 | >10pk-yr | pT2 | pN2 | 2 | Surgery + CRT | Cisplatin | >10pk-yr | pT2 | pN2 | 2 | Surgery + CRT | Cisplatin |
| 51 | >10pk-yr | pT3 | pN1 | 2 | Surgery + CRT | Cisplatin | >10pk-yr | pT3 | pN1 | 2 | Surgery + CRT | Cisplatin |
| 52* | ≤10pk-yr | pT2 | pN0 | 1 | Surgery + RT | | >10pk-yr | pT2 | pN0 | 1 | Surgery + RT | |
| 54 | ≤10pk-yr | pT2 | pN2 | 2 | Surgery + CRT | Cisplatin | ≤10pk-yr | pT2 | pN2 | 2 | Surgery + CRT | Cisplatin |

*Cases and controls were sequenced from primary tumor and not metastatic lymph nodes.

†Same control was used for two cases

pk-yr = pack years, RT = radiation therapy, CRT = chemoradiation

**Supplementary Table 4: Characteristics of HPV+ OPSCC patients in three validation cohorts**

| Cohort | Total patients | Patients receiving nonsurgical therapy | Patients with recurrence | Months to last recurrence event | Potential controls |
|---|---|---|---|---|---|
| UNC | 89 | 81 | 15 | 58.6 | 53 |
| JHU | 47 | 16 | 4 | 48.7 | 19 |
| TCGA | 53 | 29 | 6 | 41.6 | 12 |

### Supplementary Methods

#### Matching strategy in absence of a perfect control match

Six recurrent cases for which optimal pT and/or pN control matches were not identified without disrupting matching by the other criteria (R16, R24, R28, R29, R37, R40) were paired with controls with different pT and/or pN but the same pathologic overall stage. One of these six cases (R16), which received adjuvant cetuximab, was matched with a control that received cisplatin. These differences are highlighted in Supplementary Table 3.

#### Sample description, mRNA Library preparation and sequencing

Two tumor tissue sample were macrodissected from FFPE blocks of each patient. RNA sample quality was assessed by RNA TapeStation (Agilent Technologies) and quantified by AccuBlue® Broad Range RNA Quantitation assay (Biotium). Total RNA libraries were created using Illumina Stranded with RiboZero plus kit. Final library quantity was estimated by Qubit 2.0 (ThermoFisher) and quality was assessed by TapeStation HSD1000 ScreenTape (Agilent Technologies). Final library size was ~350bp with an insert size of about 200bp based on Illumina® 8-nt unique dual-indices. Libraries were pooled in equimolar proportions and sequenced on an Illumina® Novaseq platform with a read length configuration of 150 for 80M paired end reads per sample (40M in each direction).

#### Processing of RNASeq data

Low-quality bases and adapter sequences from raw RNA sequencing FASTQ files were trimmed using Trimmomatic-0.32 [1]. Filtered high quality sequencing reads were then aligned to the GRCh38 genome [2] with STAR v2.7.1a [3] with –twopassMode, --outFilterIntronMotifs and RemoveNoncanonical parameters. Quantification of reads aligned to exonic regions was conducted using STAR's –quant mode and HTSeq2.0 [4]. DeSeq2 collapse replicate was used for combining replicates and normalization [5].

#### Viral transcript alignment

Revised HPV16 reference genome and genomic annotations were obtained from the papillomavirus episteme (PaVE) [6]. Star genome index was generated using genomeSAindexNbases 8 as per size of HPV16 genome (7904 bases). Unaligned reads after human reference genome alignment were used to perform alignment with revised HPV16 viral genome with using STAR v2.7.1a –twopassMode. Quantification of reads aligned to exonic regions was conducted using STAR's –quant mode and HTSeq2.0[4].

#### GSEA, GSVA, and calculation of combined score from TPS and ISS

Gene set enrichment analysis (GSEA) [7] was performed on the Hallmark gene set (H subset, n=50 pathways) [8] and non-disease-associated KEGG Legacy pathways (subset of CP pathways, n=147) [9] using the fgsea R package [10] wi4th 1000 permutations. Gene set variation analysis (GSVA) scores were generated for unique genes within pathways associated with immune suppression (n=921) and tumor progression (n=1569) using the ssGSEA R package [11] with options set to kcdf =" gaussian," maximum group size= "2000", and mx.diff parameter= "true". GSVA scores were generated for external cohorts using gene lists obtained from case control cohort for tumor progression and immune suppression with same parameters.

Multivariate logistic regression analysis was used to construct a combined score from GSVA-derived ISS and TPS scores. Considering significant correlation between these scores, interaction term was also incorporated in the calculation of combined scores by the following equation: *Combined Score= (TPS x 2.826) + (ISS x 3.521) + (ISS x TPS x -1.894) – 2.66*.

#### Details of R v4.2 usage

Youden-index was calculated using the cutpointr R package [12]. R package pROC was used for ROC analysis [13]. For survival analysis, the Survminer [14] R package was used.

#### Immunohistochemistry details

Heat-induced epitope retrieval was performed per manufacture protocol using combinations of the following two buffers: Epitope Retrieval 1 (ER1) (Leica, Cat#AR9961) and Epitope Retrieval 2 (ER2) (Leica, Cat#AR9640). IHC staining was performed by one of two protocols, HUP 60/20 and HUP Refine, which differ in primary antibody incubation times. The HUP Refine protocol uses an incubation time of 15 minutes with the primary antibody and 8 minutes with the secondary antibody. The HUP 60/20 protocol uses an incubation time of 60 minutes for primary antibody and 20 minutes for secondary antibody. The anti-mouse or anti-rabbit secondary antibodies used in both protocols are part of the Refine Detection Kit (Cat#DS9800).
